# Effect of entomopathogenic fungus *Metarhizium robertsii* on disease incidence of Faba beans (*Vicia faba* L.) in field conditions

**DOI:** 10.1101/2022.06.11.495633

**Authors:** Lyudmila F. Ashmarina, Tatiana A. Sadokhina, Maxim Tyurin, Victor P. Danilovb, Victor Glupov

## Abstract

Faba beans (*Vicia faba* L.) are affected by a large number of pathogens. Studies carried out in western Siberia revealed the influence of the entomopathogenic fungus *Metarhizium robertsii* on reducing the development and prevalence of a complex of diseases in faba bean in the field. There was a decreasing tendency of the level of phytopathogen infection of seed material and fragments of underground organs during presowing treatment of the bean seeds with *M. robertsii*. Treatment with *M. robertsii* significantly reduced the development of root rot by 2.9 and 9.5 times in 2019 and 2020, respectively, and the prevalence of the disease decreased by 2.9-3.0 times. It also resulted in reduction of the severity of diseases of aerial organs during the growing season on average for the plant: the disease development index (DDI) of powdery mildew by 3.8 times and chocolate spot by 3.5 times. There was a significant increase in the number of active nodules on the roots of plants during the treatment with *M. robertsii*. The results obtained indicate that the treatment of bean seeds with the entomopathogenic fungus *M. robertsii* improved the phytosanitary situation of the sowing of the crop, and in the future, this technique can be used in agricultural practice.

## 1. Introduction

Entomopathogenic fungi from genera such as *Beauveria* and *Metarhizium* are able to colonize various plants, forming stable endophytic systems (Vega, 2018). These fungi can negatively affect phytophagous insects and phytopathogenic microorganisms and increase the adaptive properties of plants (Ownley *et al*. 2010; Sasan and Bidochka, 2012). Entomopathogenic fungi positively affect the structure of the soil, increase the absorption of water and nitrogen by plants, and increase the resistance of plants to abiotic and biotic environmental factors due to the production of various metabolites (amino acids, vitamins, phytohormones and antioxidant enzymes), which can lead to an increase in yield (Ownley *et al*. 2010; Busby *et al*. 2016; Hu and Bidochka, 2021). Considering the abovementioned factors, entomopathogenic fungi are promising for the creation of complex biological products for their use in integrated systems to protect plants from insects and phytopathogens. The genus *Metarhizium* plays a special role among entomopathogenic fungi. These fungi are regular representatives of soil mycobiota capable of exhibiting endophytic properties and entering mutualistic interactions with plants; they can suppress infections caused by various phytopathogens and are also widely used against agricultural pests (Vega, 2018). Despite a large number of studies showing the active colonization of plants by fungi of the genus *Metarhizium*, the ecology of the fungi remains unclear. So, in a number of stadies, it is believed that Metarhizium fungi tend to the rhizosphere zone (Vaga 2018). It is known that under initial deposit of entomopathogenic fungi in the soil plant colonization occurs extremely rarely, as it has been shown in the recent paper by Tyurin et al. (2020) on potato plants. At the same time, a number of experiments have shown a decrease in plant infection with pathogens localized in the soil (Vega 2018.; Tomilova et al. 2020).

It is quite natural that this fungus is used for plant protection, including legumes. It should be noted that legumes, including faba bean (*Vicia faba* L.), are one of the oldest crops grown in 58 countries of the world and ranked third among legumes (Sing *et al*. 2013). Various diseases significantly affect the yield of legumes. The value of losses of the production of legumes caused by diseases is estimated at an average of 15%, sometimes reaching 70-80% (Horoszkiewicz-Janka *et al*. 2013). Diseases of legumes can be caused by representatives of various taxonomic orders: fungi, viruses, and bacteria (Mohammed, 2013).

Under the conditions of the continental climate of western Siberia, a whole range of diseases is widespread in crops of faba beans: root rot (*Fusarium* L., *Alternaria* L.), leaf spots (*Fusarium, Alternaria*), powdery mildew (*Erysiphe communis* (Wallr.) Grev. f. faba Yacz.), and chocolate spot (*Botrytis fabae* Sardina) (Ashmarina *et al*. 2010). Strong manifestations of diseases and their widespread presence significantly reduce the productivity of legumes, which results in the use of chemical methods of protection and, therefore, environmental pollution (Ashmarina *et al*. 2017).

Unlike most chemicals that are active only against insects or plant pathogens, fungi of the *Metarhizium* genus can have a complex effect on both insect pests and plant pathogens (Vega, 2018). The ecology of these species in agroecosystems in temperate regions, such as Europe and North America, has been well studied (Meyling and Eilenberg, 2007). In the extreme conditions of western Siberia (Russia), there are only a few studies on the effect of the entomopathogenic fungus *Metarhizium robertsii* on plants in the field (Tomilova *et al*. 2020). In addition, most studies on the antagonistic effects of *Metarhizium* species have been carried out on one or more phytopathogens and mainly in laboratory studies (in vivo or in vitro), that is, “with limited external validity” (Vega, 2018). According to Barra-Bucareli *et al*., 2019, while this provides important scientific validity, it does not necessarily guarantee similar performance under field conditions. The purpose of our research was to evaluate the effect of *M. robertsii* on the complex of the main diseases that occur in *V. faba* cultivated in the field under continental conditions in western Siberia.

## 2. Materials and methods

### 2.1 Preparing fungi material

We used isolates of the entomopathogenic fungus *M. robertsii* (isolate P-72) and from the collection of microorganisms of the Institute of Systematics and Ecology of Animals (Siberian Branch of the Russian Academy of Science). The fungal species were identified on the basis of the sequence of the translation elongation factor (EF1 α) region (Kryukov *et al*. 2017).

The mycelium was grown by two-phase cultivation. First, a deep culture of the fungi was grown in Sabouraud’s dextrose agar (SDA) in an incubator shaker at 160 rpm at 26°C for 4 days. Second, the fungi were cultivated on twice-autoclaved millet. Then, millet with conidia was dried for 10 d at 24 °C and 27% RH and homogenized with a ball mill.

### 2.2 Field experiment

The field studies were carried out in 2019 and 2020 at the station of the Siberian Research Institute of Forage of the SFNCA RAS located in the northern forest-steppe of the Ob region of the Novosibirsk region of Russia. The soil type was leached chernozem, medium thick, medium loamy, and the organic carbon content in the soil was 3.48% and pH was 5.3. The amount of absorbed bases was 58–61 mg/eq. per 100 g of soil. Before crop rotation the field was sowing in pairs. There were natural initial deposit of root rot pathogens in the soil. Root rot pathogens were not introduced into the soil.The seeds of faba beans of the Sibirskie variety were strongly infected with root rot pathogens were used for sowing: in 2019, the level of fungi of the genus *Fusarium* was 12.0% and that of the genus *Alternaria* was 12.0%; in 2020, *Alternaria* was 57.0% and Fusarium was 8.0%.

Fungal conidia were suspended in a water-Tween solution (Tween 20, 0.04%). Faba bean seeds of the experimental variant were inoculated with a suspension of *M. robertsii* conidia with a titer of 5 × 107 conidia/mL (2.5 L per 20 kg of seeds) and left to dry at a temperature of 18-20 °C. The control variant in the experiment was treated with an aqueous solution of Tween 20. The placement of variants in the experiment was systematic; in 2019, there was fourfold replication, and in 2020, there was fivefold replication. Sowing of faba bean was carried out in 2019 on May 16 and in 2020 on May 19, when the soil temperature at a depth of 6-8 cm reached 8-10°C. Sowing was carried out with an Optima planter (Kverneland Group Soest, GmbH). The seeding depth was 6-8 cm. The seeding rate was 400 thousand germinating grains per hectare. The plot length was 10 m, the width was 3.9 m, and the accounting plot area was 39 m2. Harvesting was carried out with a Wintersteiger Classic grain harvester (Wintersteiger AG, Austria).

### 2.2 Experimental time points

After treating the seeds of the beans with *M. robertsii*, samples of the seed material were collected to analyse the effectiveness of their action on phytopathogens. In the field, we manually excavated and assessed the damage to the bean plants by root rot 5 weeks after sowing in 2019 and 4 weeks after sowing in 2020.

Plants developed differently during the research. In 2019, the growing season was extended, and the phases of plant development lagged by a week from similar phases in 2020. Observations in the field for the damage to plants by spotting were carried out in the phases of stem elongation and flower initiation in 2019 on days 35 and 57 after sowing and in 2020 on days 28 and 50. Diseases of the aboveground parts of plants (powdery mildew, fusarium, chocolate spot, and others) were counted on day 80. For mycological analysis, plants were selected on the 35^th^ day after sowing in 2019 and on the 28^th^ day in 2020. The characteristics of plant growth and development were evaluated on days 12, 26, 41, 59, 79 and 98. For this, 10 plants were randomly excavated from each replication (n = 50 plants in each variant). The following phases of development were recorded in beans: full shoots, 5-6 leaves, branching, flowering, bean formation, seed filling and maturation. Productivity was analysed by evaluating ten random plants in each replication at harvest time. The weight of 1000 grains and the number and weight of grains from each plant were estimated (Schwerz *et al*. 2019).

The water concentration in the soil was measured layer by layer every 10 cm to a depth of 1 m with an Aquater M-350 (STEP Systems GmbH (Germany)).

### 2.3 Weather conditions during faba bean growth

The 2019-2020 growing season was characterized as close to the climatic norm (hydrothermal coefficient (HTC)). In 2019, the HTC for May-September was 1.2, with high rainfall in May and July (116% and 161% mean annual precipitation, respectively) and a lack of moisture in June (HTC – 0.5) and August (HTC – 0.4). In 2020, the HTC for May-September was 1.29, but with monthly variable amounts of precipitation and a lack of moisture in June (HTC - 0.4) and in the second week of July (HTC - 0.6) (Fig. 1). During the growing season of 2019, 263 mm of precipitation fell (corresponds to the average annual indicator, the average temperature is 15.2 ^0^C. In 2020, 310 mm of precipitation fell during the period from May to September, which is 50 mm higher than the average annual norm, the average temperature is 16.3^0^ C.

**Figure 1.**
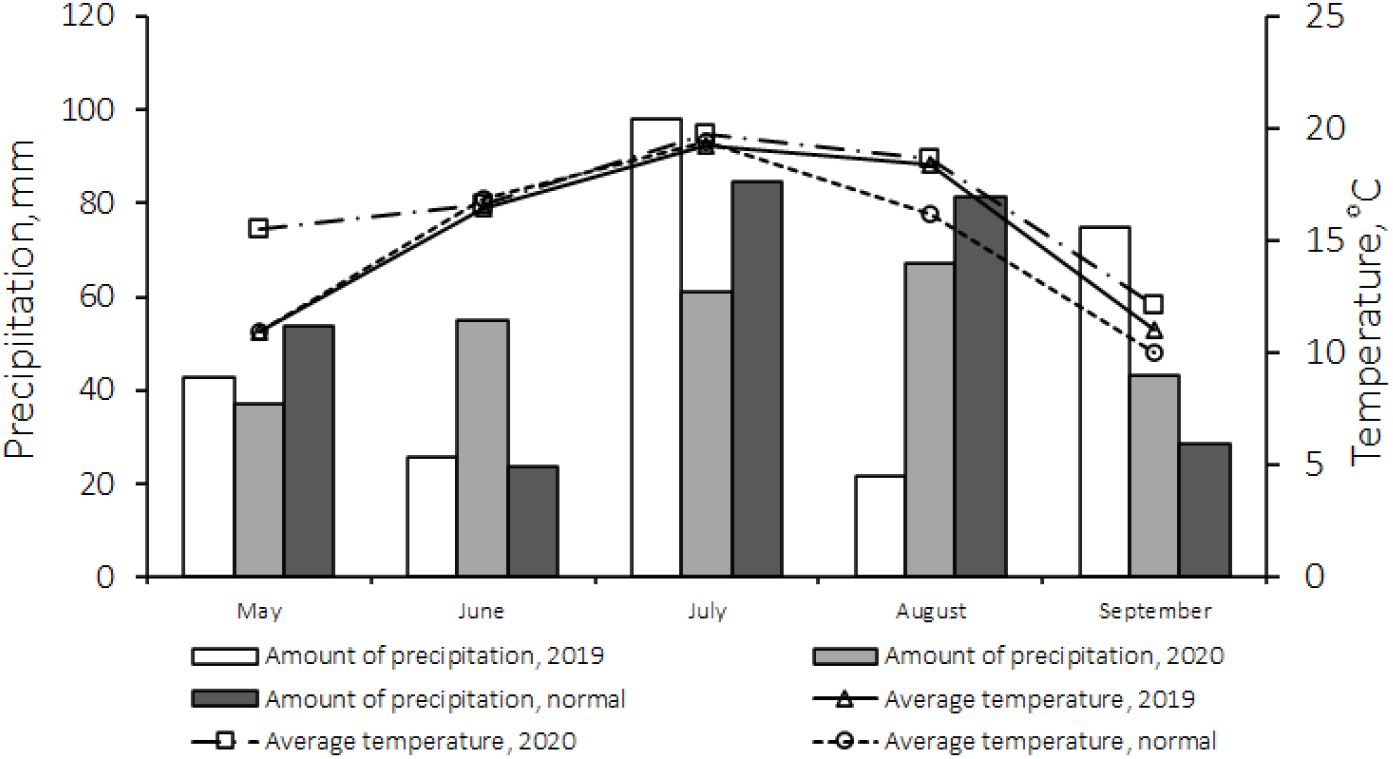
The dynamics of temperature and precipitation by month in the 2019-2020 and comparison with the average long-term data

A good supply of soil moisture in May 2019 (103 mm) contributed to the emergence of seedlings. Rainfall and temperature in June were 47% and 0.5 °C below the mean annual rate, respectively.

### 2.4 Analysis of infection of the plants

Ten plants were randomly collected from each replication (n = 50 in each variant) to determine the degree of damage and spread of diseases. The plants were excavated, freed from soil, thoroughly washed with running water and used for analysis. Analysis of the degree of damage and spread of root rot was carried out on fresh, nondried samples when the pigmentation of the affected tissue appeared brighter. The degree of damage to underground organs (root collar of the stem and roots) was determined visually using a modified 4-point scale according to Noronha et al. (1995). In particular, 0 points indicate a healthy plant; 1 point indicates mild damage to an organ or plant; 2 points indicate moderate damage without serious damage to organs or plants; 3 points indicate severe damage to some organs or plants and 4 points indicate serious organ damage and plant death. The disaese development index (DDI) damage to beans was calculated according to the following generally accepted formula: R = ∑ a × b × 100: N × K, where R is the development of the disease (points or percentages); ∑ a × b is the sum of the products of the number of diseased plants (a) by the corresponding score or percentage of damage (b); N is the number of considered plants in the sample; and K is the highest point on the scale.

### 2.5 Microbiological Analysis (in vitro)

To assess the infection of plant parts with root rot pathogens, a part of the stem adjacent to the root was isolated (Jensen *et al*. 2020). The resulting root fragments 1 cm in length were superficially sterilized as described previously. They were then placed on sterile Czapek medium supplemented with streptomycin (0.6 g/L) and incubated in a thermostat (24 °C) for 14 days (Miché and Balandreau, 2001).

Phytopathogen identification was carried out by light microscopy (Gerlach and Nirenberg, 1982; Seifert and Gams, 2011). Similar actions were carried out during seed analysis.

The severity of damage to leaf-stem infections was considered using the recommendations set out in previous papers (Ashmarina *et al*. 2010). The development of the disease was calculated according to the generally accepted formula.

The biological effectiveness (BE) of the use of entomopathogenic fungi was calculated using the formula BE = (R - r)/R x 100%, where R is the index of damage in the control and r is the index of damage in the variants with *M. robertsii* treatment.

To count rhizobial nodules in beans, 10 plants were randomly selected from each repetition (n = 50) on days 41, 59, and 79 after sowing. Then, the root system was thoroughly washed under running water, and the number of nodules on each plant was counted.

### 2.6 Microbiological Analysis of Plant Colonization (in vitro)

Colonization of plants and soil by *M. robertsii* entomopathogenic fungi was assessed by plating of plant particles or aliquots of soil solution on modified nutrient medium (glucose 40 g/L peptone 10 g/L, cetyltrimethylammonium bromide 0.35 g/L, cycloheximide 0.05 g/L, tetracycline 0.05 g/L, and streptomycin 0.6 g/L) in 90 mm Petri dishes. The dishes were incubated at 24 °C for 14 days, and *Metarhizium* colonies were detected by light microscopy and counted. A weighed portion of each soil sample was dried at 60 °C for 24 h, and the colony-forming unit (CFU) count was adjusted to the dry weight of the soils.

To analyse the colonization of plants by entomopathogenic fungi, the roots, stems and leaves were examined (5 plants from each replication and 20 plants from each variant). To assess endophytic colonization by the entomopathogenic fungi, we selected the middle part of the root, the lower third of the stem, and the leaf from the middle plant layer. The plant parts were washed with running water and sterilized with 0.5% sodium hypochlorite and 70% ethanol (Posada *et al*. 2007). The organs were imprinted on the abovementioned medium and then placed on the surface of the medium in 90 mm Petri dishes (McKinnon *et al*. 2017). After 14–20 days of incubation, the growth of *Metarhizium* was detected visually and by light microscopy. The percentage of fungus-positive plants was then calculated. Samples showing fungal growth on the prints were excluded from analysis. To analyse colonization of the nonsterilized roots, the middle parts of the roots were washed 3 times (1 min at 180 rpm each time) in a water–Tween 20 solution (0.04%) and plated on the abovementioned medium in petri dishes. Incubation and detection of the fungi were performed as described above.

To assess the number of CFUs in bulk soils, 5 g of sample was suspended in 40 ml of a sterile water-Tween solution (0.1%), vortexed for 10 s, and shaken at 180 rpm for one hour. A 100 μL aliquot of the soil suspension from each sample was plated on the medium in 90 mm Petri dishes. The incubation and detection of fungi were performed as described above.

### 2.7 Statistical analysis

Data analysis was performed using Statistica 8 (StatSoft, Inc., USA) and PAST 3 (Hammer, *et al*. 2001). The normality of the data distribution was checked using the Shapiro– Wilk W test. Normally distributed data were analysed by one-way analysis of variance followed by a post hoc LSD Fisher test. Abnormal data were analysed using the Kruskal– Wallis test and then the posterior Mann–Whitney test. Student’s t test was used to assess the number of rhizobial nodules. Fisher’s exact test was used to assess differences in plant colonization by fungi.

## 3. Results

### 3.1 Effect of treatment of faba bean seeds with *M. robertsii* on the development of pathogens before sowing (in vitro)

The year 2020 was characterized by a higher level of infection with root rot pathogens. A steady tendency of reduction of the infection of bean seeds with root rot pathogens when treating seeds with *M. robertsii* was established. Thus, in comparison with the control, the infection of seeds by species of the genus *Fusarium* in 2019-2020 decreased by 3.8-3.1 times (Table 1). In 2020, *M. robertsii* reliably suppressed only fungi of the genus *Alternaria* (Fisher, p = 0.00001) on seeds, while the biological efficiency was 63.3%. Although in relation to species of the genera *Fusarium* and *Cladosporium*, the biological efficiency in 2020 was high (89.3 and 73.7%, respectively), no statistically significant differences were found. It should also be noted that seed treatment with *M. robertsii* significantly reduced the infection of seeds with fungi of the genus *Aspergillus* (Table 1).

**Table 1.**
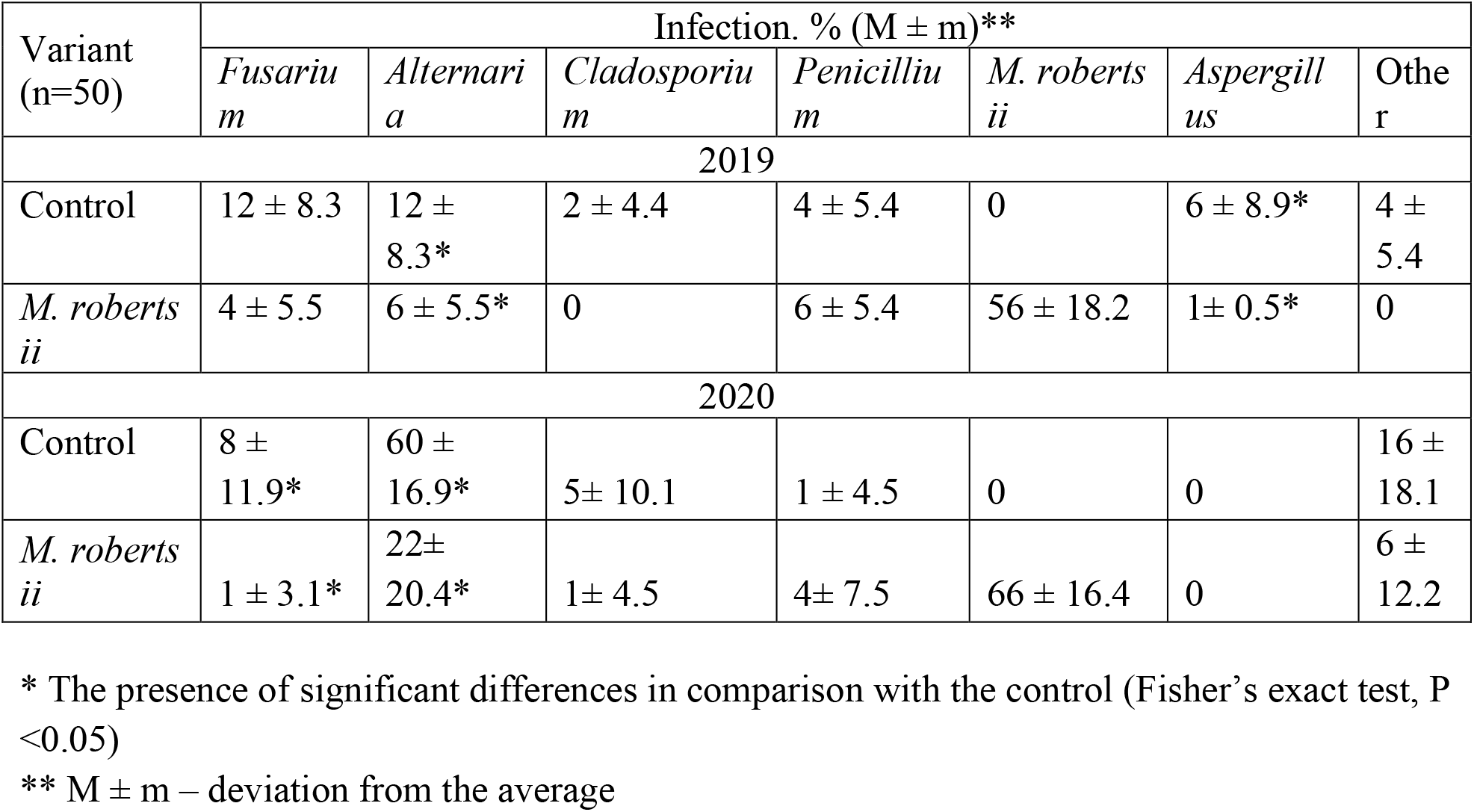
Influence of pre-sowing treatment of seeds of *M. robertsii* beans on their infection

| Variant<br>(n=50) | Infection. % (M ± m)** |  |  |  |  |  |  |
| --- | --- | --- | --- | --- | --- | --- | --- |
|  | *Fusarium* | *Alternaria* | *Cladosporium* | *Penicillium* | *M. robertsii* | *Aspergillus* | Other |
| 2019 |  |  |  |  |  |  |  |
| Control | 12 ± 8.3 | 12 ± 8.3* | 2 ± 4.4 | 4 ± 5.4 | 0 | 6 ± 8.9* | 4 ± 5.4 |
| *M. robertsii* | 4 ± 5.5 | 6 ± 5.5* | 0 | 6 ± 5.4 | 56 ± 18.2 | 1 ± 0.5* | 0 |
| 2020 |  |  |  |  |  |  |  |
| Control | 8 ± 11.9* | 60 ± 16.9* | 5 ± 10.1 | 1 ± 4.5 | 0 | 0 | 16 ± 18.1 |
| *M. robertsii* | 1 ± 3.1* | 22 ± 20.4* | 1 ± 4.5 | 4 ± 7.5 | 66 ± 16.4 | 0 | 6 ± 12.2 |

\* The presence of significant differences in comparison with the control (Fisher's exact test, P < 0.05)
\*\* M ± m – deviation from the average

### 3.2 Influence of *M. robertsii* treatment on the development of root rot during the growing season of the plants

Root rot of beans was mainly represented by species of the genera *Fusarium* spp., *Alternaria* spp., and *Cladosporium* spp. Significant infection of the seeds in 2020 led to a higher development of the disease (4.5 times higher) than that in 2019. It was shown that *M. robertsii* actively suppressed the development of root rot in the early (4 weeks after sowing) stages of plant growth (Fig. 2A). Significant differences in disease development between plants treated with *M. robertsii* and controls were established. In plants treated with *M. robertsii*, the damage in 2019 was 2.9 times lower than that in the control (Mann–Whitney test, p = 0.025) and 9.5 times lower in 2020 (Man-W, p = 0.002). When plants were treated with *M. robertsii*, the prevalence of root rot also decreased (Fig. 2B). In 2019, the prevalence of the disease was 2.9 times lower when processing plants with *M. robertsii* (Mann–Whitney test, p = 0.007, compared with the control) and 3.0 times lower in 2020 (Mann–Whitney test, p = 0.025, compared with the control). The biological effectiveness in reducing the prevalence of disease during seed treatment in 2019 and 2020 was 65.4% and 66.7%, respectively.

**Figure 2.**
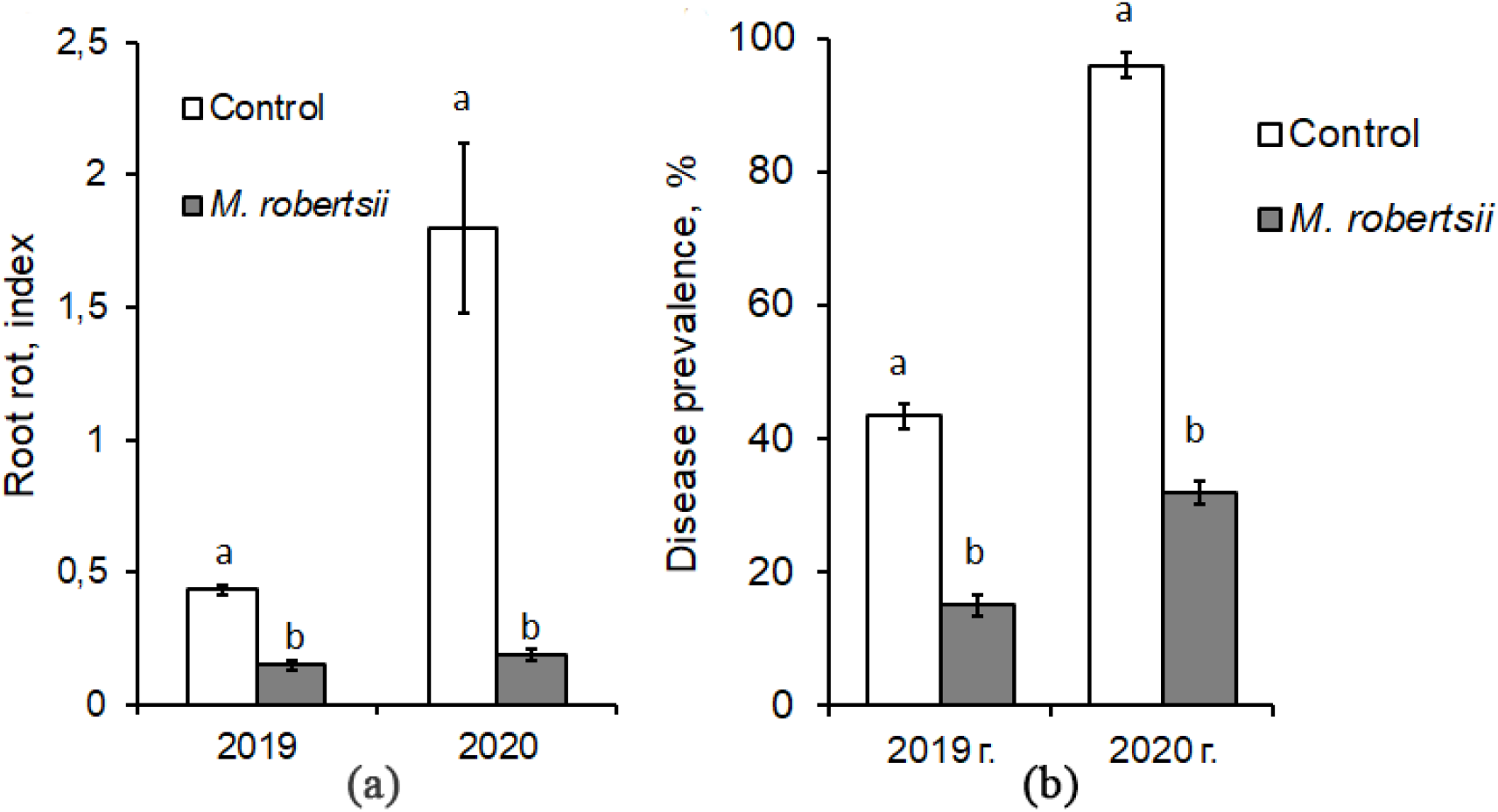
Effect of the treatment of seed beans with entomopathogenic fungi during the 2019-2020 (a) damage to beans caused by root rot disease expressed as root rot developmen index; (b) level prevalence root rot disease on plant of bean. The vertical lines indicate the standard error of the averages. The same letters indicate insignificant differences between all treatments (Mann–Whitney test, P **<**0.05)

Plants treated with *M. robertsii* also had a more powerful root system, an increase in the number of lateral roots and the formation of active nitrogen-fixing rhizobial nodules (Table 2)

**Table 2.** The number of active nodules on the roots of *Vicia faba*, 2019-2020

| Variant<br>(n=50) | Number of active nodules, pieces / plant |  |  |
| --- | --- | --- | --- |
|  | 36 days after sowing | 55 days after sowing | 73 days after sowing |
| 2019 |  |  |  |
| Control | $6.4 \pm 1.559$ | $10.4 \pm 5.7881$ | $5.8 \pm 2.6073$ |
| *M. robertsii* | $7.3 \pm 2.854$ | $12.3 \pm 6.786$ | $8.1 \pm 2.8771^*$ |
| 2020 |  |  |  |
|  | 41 days after sowing | 59 days after sowing | 79 days after sowing |
| Control | $1.6 \pm 0.7495$ | $6.8 \pm 2.3486$ | $3.2 \pm 1.3246$ |
| *M. robertsii* | $3.8 \pm 1.4538$ | $20.3 \pm 8.3432^*$ | $4.7 \pm 1.9845$ |

\* The presence of significant differences in comparison with the control (Fisher's exact test, $P < 0.05$ ).

In 2019, at the beginning of the growing season, an insignificant increase in the number of nodules was noted when processing beans compared with the control (Mann– Whitney test, p = 0.3680). By the end of the growing season (73 days after sowing), the difference from the control increased by 1.4 times (Mann–Whitney test, p = 0.0407). In 2020, 41 days after sowing, the number of active nodules on the roots of beans in the control variant was 2.4 times less (T Test, p = 0.0001) than in the treated plants. The greatest differences from the control were noted 8 weeks after seeding - by 3.0 times (T Test, p = 0.0002). By the end of the growing season (11 weeks after sowing), the discrepancy levelled off (T test, p = 0.0007). A slight development of rhizobial nodules on the 79th day was associated with a drought that developed at the end of June 2020 - GTC 0.4, and the moisture reserve in the soil in the 0-20 cm layer was 10.5 mm. Mycological analysis of sections of underground organs of bean plants collected 4 weeks after germination revealed a tendency to improve the phytosanitary condition when treated with *M. robertsii* (Table 3). Mycological analysis of sections of the underground organs of bean plants collected 4 weeks after germination revealed a tendency for their recovery when treated with *M. robertsii* (Table 3). Compared with the control in 2019, in the variants treated with the entomopathogenic fungus *M. robertsii*, there was a trend towards a decrease in the colonies of fungi of *Fusarium* by 3.2 times, of *Alternaria* by 2.3 times, and species of *Cladosporium* by 2.0 times. The 2020 data confirmed the results obtained on the reduction of plant infestation by root rot pathogens in the treated variants. The number of colonies of fungi of the genus *Alternaria* significantly decreased by 4.0 times (Fisher, p = 0.113) and of the genus *Fusarium* by 1.3 times (Fisher, p = 0.591). It should be noted that a significant infection of plants by *Fusarium* species led to damage to the vascular system and wilt of the plants.

**Table 3.** Infection of beans with root rot pathogens, 2019-2020

| Variant | Number of colonies in a Petri dish ($n = 5$, $M \pm m^{**}$) | | | | | |
| --- | --- | --- | --- | --- | --- | --- |
|  | *Fusarium* | *Alternaria* | *Cladosporium* | *Penicillium* | *Aspergillum* | *Other* |
| 2019 |  |  |  |  |  |  |
| Control | $1.6 \pm 0.4$ | $1.6 \pm 0.4$ | $1.0 \pm 0.55$ | $0.6 \pm 0.24$ | $0.8 \pm 0.2$ | $2.0 \pm 0.71$ |
| *M. robertsii* | $0.5 \pm 0.22$ | $0.67 \pm 0.33$ | $0.5 \pm 0.22$ | $0.5 \pm 0.22$ | $0.33 \pm 0.21$ | $1.0 \pm 0.26$ |
| 2020 |  |  |  |  |  |  |
| Control | $8.0 \pm 0.55$ | $2.4 \pm 0.81^*$ | $1.0 \pm 0.32$ | 0 | 0 | $0.4 \pm 0.24$ |
| *M. robertsii* | $6.4 \pm 0.51$ | $0.6 \pm 0.24^*$ | $0.4 \pm 0.24$ | 0 | 0 | 0 |

\* The presence of significant differences in comparison with the control (Fisher's exact test, $P < 0.05$ )
\*\* $M \pm m$ – deviation from the average

### 3.3 Dynamics of the development of diseases on the aboveground organs of the beans

It is known that severe leaf damage leads to a decrease in photosynthesis, enzymatic processes and other physiological functions and affects the productivity of plants.

It was found that the entomopathogenic fungus *M. robertsii* actively suppressed the development of spots on plants (Fig. 3). Therefore, in 2020, the level of disease manifestation was significantly lower than that in 2019, which was primarily due to weather conditions favourable for the development of phytopathogens.

**Figure 3.**
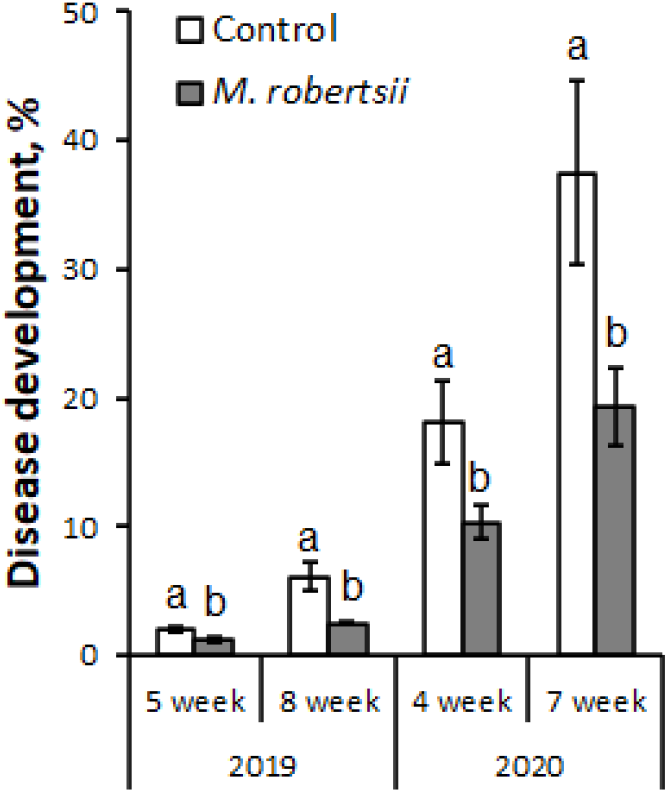
Effect of seed treatment of M. robertsii bean seeds on the development of spotting in 2019 and 2020. The vertical lines indicate standard error of the means (4 biological replicates, ten plants each). The same letters indicate significant differences in DDI between control and M. robertsii treatment (Man-W, P<0.05)

The dry weather of 2019 restrained the manifestation of the disease. In the first and second weeks of July, a large amount of precipitation fell (200% and 337% of the average annual precipitation, respectively), which favorably affected the development of plantsThe acutely arid period in the third week of July (2.0 mm of precipitation) and the first week of August (1.6 mm of precipitation) held back the development of leaf-stem infections. The end of August and the beginning of September were characterized by high temperatures and moderate rainfall. This weather favoured a longer growing season for the bean plants. In general, the weather conditions in 2019 were favourable for the development of legume plants and not favourable for the development of phytopathogens.

The hydrothermal conditions of the 2020 growing season significantly differed from the mean annual data. There were higher temperatures in May (on average 4.6°C higher), temperatures close to normal in July and June, and an uneven distribution of precipitation.

During all months of the growing season (except June), the average annual precipitation was exceeded (on average 145% of the norm). There was a deficit in June (23.8 mm if 43% of the norm).

During the sowing period of the faba beans (third week of May), weather conditions were favourable for both the growth and development of plants and the development of phytopathogens and entomopathogenic fungi. Warm and humid May weather favoured the development of plant leaf spots. The second and third weeks of June were characterized by a lack of precipitation and high temperatures (air temperature was 1.1 °C higher than the mean annual norm). These conditions had a stressful effect on the growth and development of bean plants and reduced the spread rate of diseases. A significant amount of precipitation fell in the first and third weeks of July (198 and 139% of the norm, respectively). Warm and humid weather contributed to the rapid and significant development of diseases on the leaves: powdery mildew and chocolate spot. The second half of the growing season was characterized by increased precipitation and moderate air temperatures, which contributed to the epiphytotic manifestation of the diseases.

In 2019, 5 weeks after sowing, the disease mildly manifested in the lower layer of plants, which was facilitated by the dry weather of the first week of June. In the variants treated with *M. robertsii*, when compared with untreated plants, we recorded a significant decrease in the development of spots by 1.8 times (Mann–Whitney test, p = 0.026) and 2.4 times 8 weeks after sowing (Mann–Whitney test, p = 0.028) (Fig. 3).

Similar data were obtained in 2020; 4 weeks after sowing, the disease development index (DDI) of the control variants was significantly higher (1.8 times) than that of plants treated with *M. robertsii* (Mann–Whitney test, p = 0.03). By the flowering phase (7 weeks after sowing), the disease level in the control increased to 37.5%, while in the treated variants, it did not exceed 23% (Mann–Whitney test, p = 0.029) (Fig. 3). We recorded the greatest manifestation of diseases 11 weeks after sowing during the seed-filling phase (Fig. 4).

**Figure 4.**
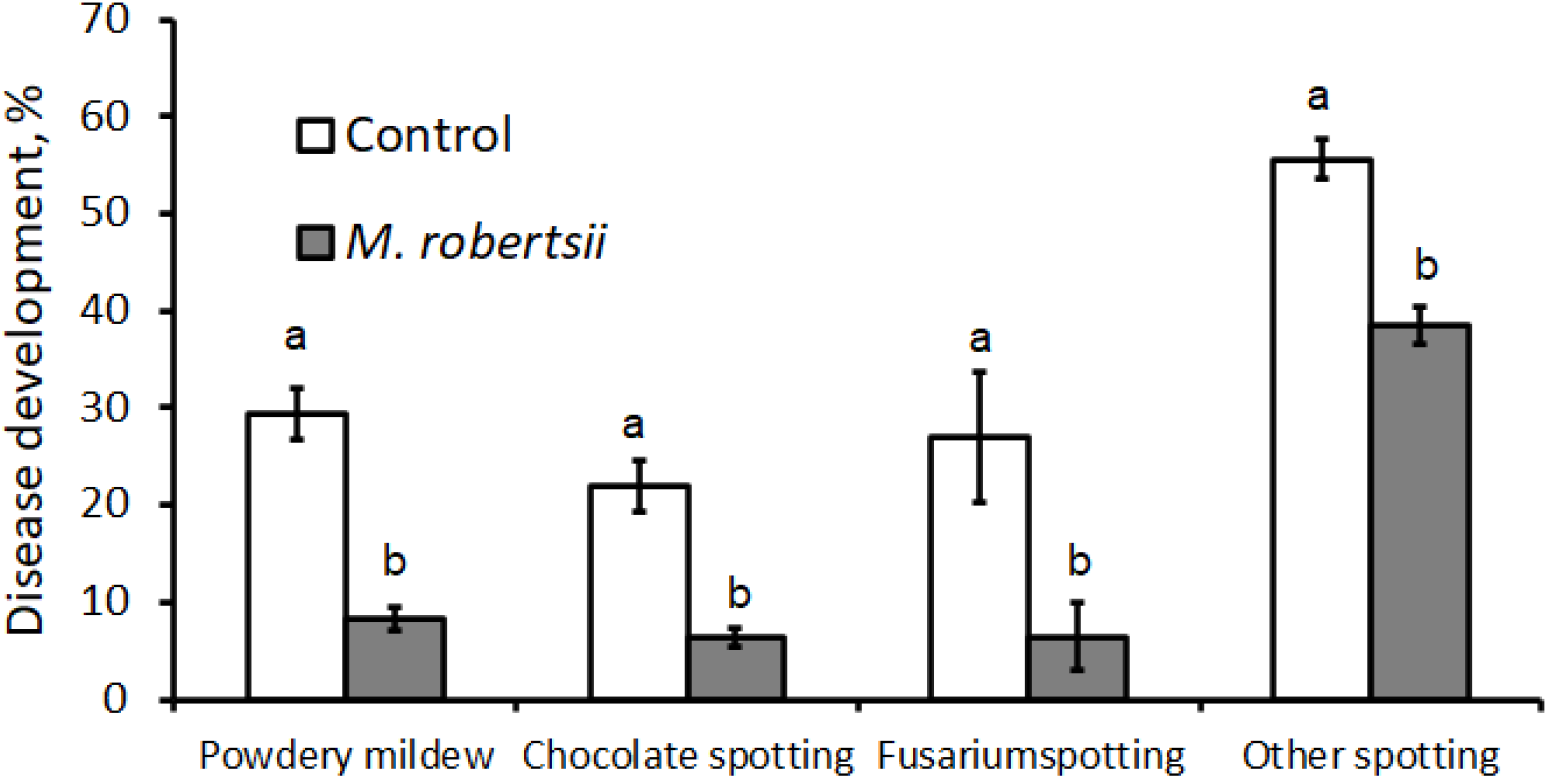
Development of various diseases on bean plants (average per plant) 11 weeks after sowing. The vertical lines indicate the standard error of the averages. The same letters indicate insignificant differences between all treatments (Mann–Whitney test, P <0,05)

Eleven weeks after sowing, there was a decrease in the DDI in all layers of *M. robertsii*-treated bean plants. When assessing powdery mildew caused by *E. communis* on crops, the DDI of the upper and middle layers of plants was 3.4-4.1 times lower than that of untreated variants (Table 4). The disease was practically absent on the lower leaves. The level of development of chocolate spot caused by *B. fabae* had a stable tendency of a healing effect when using *M. robertsii*. The DDI of the upper layer of leaves decreased 4.0 times (Mann– Whitney test, p = 0.029) and, on average, 2.8 times (Mann–Whitney test, p = 0.031). In the lower layer of the plants, a tendency towards suppression of the disease was observed, but no differences were found compared with the control (Mann–Whitney test, p = 0.058). The development of *Fusarium* disease on the top layer differed in the control and *M. robertsii* treatments (by 4.0 times), but it was not statistically significant (Mann–Whitney test, p = 0.082).

**Table 4.** Development of diseases in all layers of M. robertsii-treated bean plants eleven weeks after sowing of 2020, expressed in %

| Variant | Powdery mildew |  |  | Fusarium spotting |  |  | Chocolate spotting |  |  |
| --- | --- | --- | --- | --- | --- | --- | --- | --- | --- |
|  | lower tier | middle tier | top layer | lower tier | middle tier | top layer | lower tier | middle tier | top layer |
| Control | 0 | 27.83 | 60.17 | 30.17 | 24.83 | 26.03 | 1.67 | 21.33 | 43 |
| *M. robertsii* | 0 | 6.8* | 17.87* | 6.37* | 6.5 | 6.5 | 0.5 | 7.57* | 10.73* |

\*– indicate significant differences (Mann–Whitney test, $P < 0.05$ )

### 3.4 Colonization of *M. robertsii* in the plant and rhizosphere zones

Along with the phytosanitary effect, we found that the treatment of faba bean seeds with the entomopathogenic fungus *M. robertsii* stimulated plant growth and increased productivity.

In 2019, when studying the amount of entomopathogenic fungi in the soil, there were no significant differences between *Metarhizium* variants and the control. However, it should be noted that the CFU of fungi in the variant with seeds treated with *Metarhizium* at the beginning and middle of the season was higher and reached 1924 ± 665 colonies/gram of dry soil, and at the end of the growing season (14 weeks after sowing), it decreased by 2-4 times. Moreover, in 2020, at the beginning of the growing season, there was a significantly higher (Mann–Whitney test, p < 0.05) occurrence of entomopathogenic fungi in the soil in the treated variants. Thus, we observed a twofold increase in the CFU in the variants treated with *Metarhizium*, followed by a decrease in August to the level of the control (Fig. 5).

**Figure 5.**
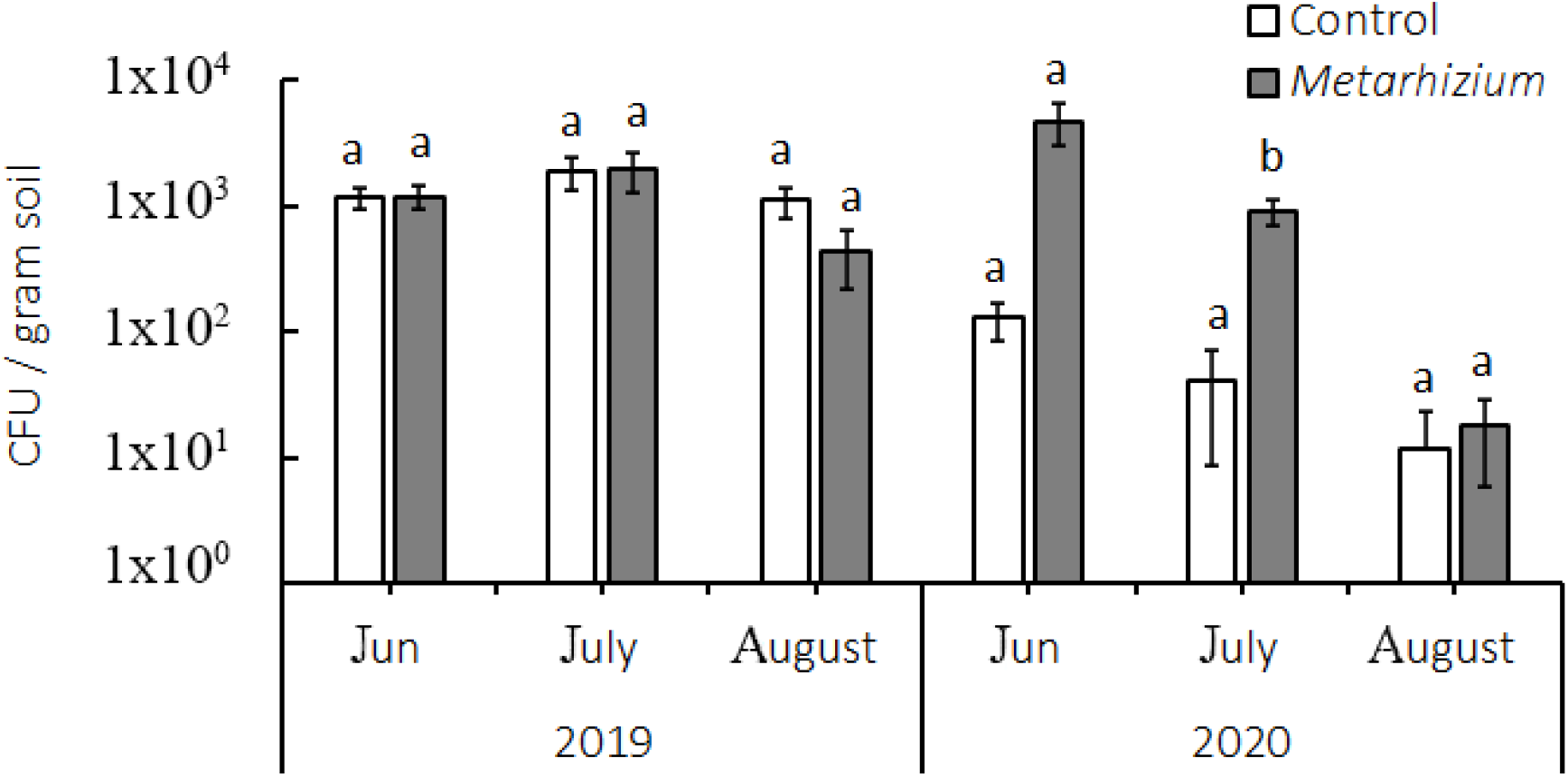
The CFU of Metarhizium in soil samples in 2019 and 2020. Vertical lines indicate the standard errors (SE). Different letters indicate significant differences as calculated for Metarhizium and each year separately (Mann-Whitney U Test, P < 0.05).

In 2019, we did not observe colonization of the tissues of the study plants (leaves, stems, and roots) by the fungus *M. robertsii*. However, in June 2020, the colonization of plant tissues by fungi in the treated variants reached 4% to 18% (*Metarhizium*-positive plants). In the rhizosphere zone (nonsterile roots), 36% *Metarhizium*-positive plants differed from the control plants, where the amount of isolated fungi did not exceed 4%. In the following months (July-August), the level of colonization decreased to the control values (Table 5).

**Table 5.**
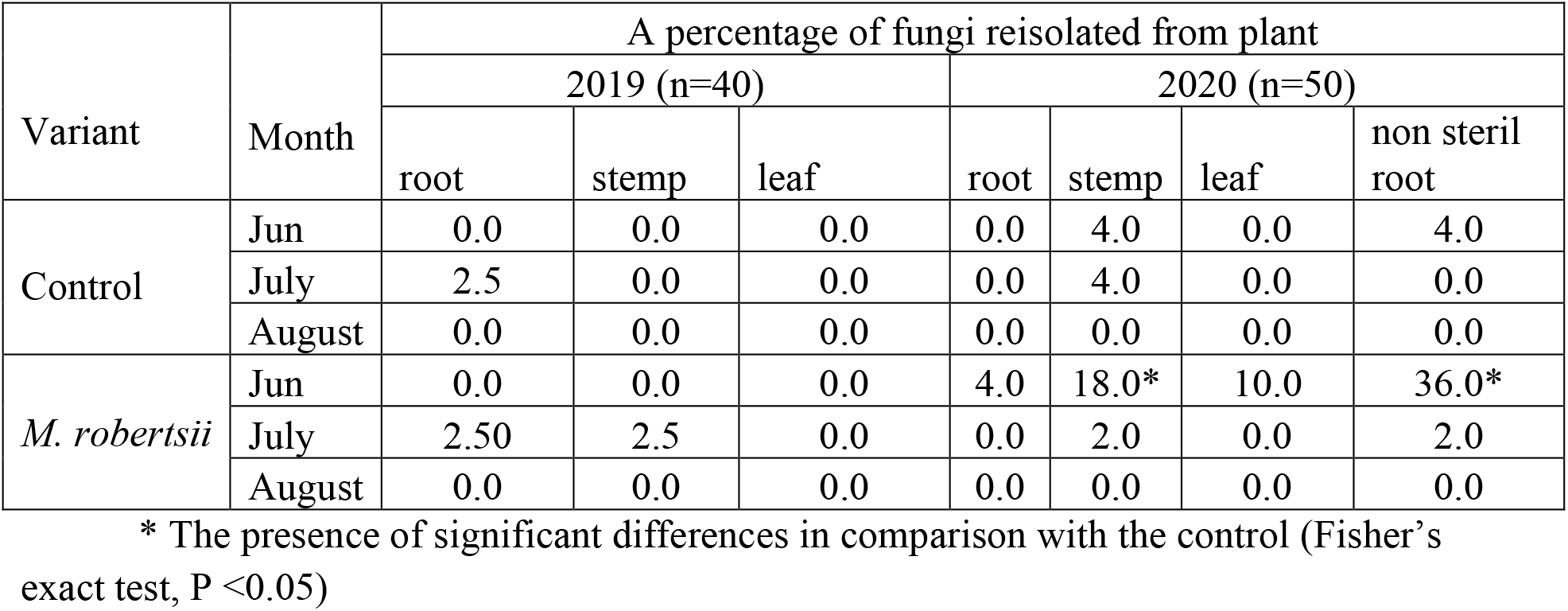
Isolation of *Metarhizium* from sterile roots, leaves and stems from plots with different types of treatment seed.

| Variant | Month | A percentage of fungi reisolated from plant |  |  |  |  |  |  |
| --- | --- | --- | --- | --- | --- | --- | --- | --- |
|  |  | 2019 (n=40) |  |  | 2020 (n=50) |  |  |  |
|  |  | root | stemp | leaf | root | stemp | leaf | non steril root |
| Control | Jun | 0.0 | 0.0 | 0.0 | 0.0 | 4.0 | 0.0 | 4.0 |
|  | July | 2.5 | 0.0 | 0.0 | 0.0 | 4.0 | 0.0 | 0.0 |
|  | August | 0.0 | 0.0 | 0.0 | 0.0 | 0.0 | 0.0 | 0.0 |
| *M. robertsii* | Jun | 0.0 | 0.0 | 0.0 | 4.0 | 18.0* | 10.0 | 36.0* |
|  | July | 2.50 | 2.5 | 0.0 | 0.0 | 2.0 | 0.0 | 2.0 |
|  | August | 0.0 | 0.0 | 0.0 | 0.0 | 0.0 | 0.0 | 0.0 |

\* The presence of significant differences in comparison with the control (Fisher's exact test, $P < 0.05$ )

### 3.5 Influence of *M. robertsii* treatment on the growth rates and productivity of beans

In 2019-2020, a significant increase in plant height was established (Man-W, p = 0.01597) in the variants treated with *M. robertsii*, which is reflected in Table 6. In general, the tendencies towards an increase in growth persisted throughout the growing season, and the strongest increase of growth by 15% (Man-W, p = 0.01219) was found in 2020, 25 days after sowing.

**Table 6.** Effect of M. robertsii bean seed treatment on plant height, expressed in cm

| Variant | Plant development (growth stage) |  |  |  |  |  |  |  |  |  |
| --- | --- | --- | --- | --- | --- | --- | --- | --- | --- | --- |
|  | 5-6 leaves |  | stem branching |  | flowering |  | bean formation |  | seed ripeness |  |
|  | 2019 | 2020 | 2019 | 2020 | 2019 | 2020 | 2019 | 2020 | 2019 | 2020 |
| Control | 15.5±<br>3.50 | 18.7±<br>0.37 | 24.4±<br>1.42 | 26.5±<br>2.69 | 38.0±<br>4.90 | 50.9±<br>2.16 | 61.8±<br>2.81 | 62.7±<br>1.44 | 78.9±<br>9.4 | 68.9±<br>2.03 |
| *M. robertsii* | 19.5±<br>2.93 | 22.0±<br>0.40 | 29.7±<br>0.89* | 30.8±<br>4.65* | 40.3±<br>3.81 | 49.6±<br>0.94 | 67.1±<br>2.16* | 68.7±<br>1.40* | 89.6±<br>9.8* | 77.5±<br>0.40* |

\*— indicate significant differences (Mann–Whitney test, $P < 0.05$ )

The yield of treated crops was higher than that of untreated plants by 17.0% (Man-W, p = 0.03038) in 2019 and by 11% in 2020 (Man-W, p = 0.03671). Along with this, the positive influence of *M. robertsii* was reflected in the increase in the mass of 1000 grains (Table 7). It was higher than the control: in 2019 by 14.8-16.2 (Man-W, p = 0.03038) and in 2020 by 8-10% (Man-W, p = 0.0367).

**Table 7.**
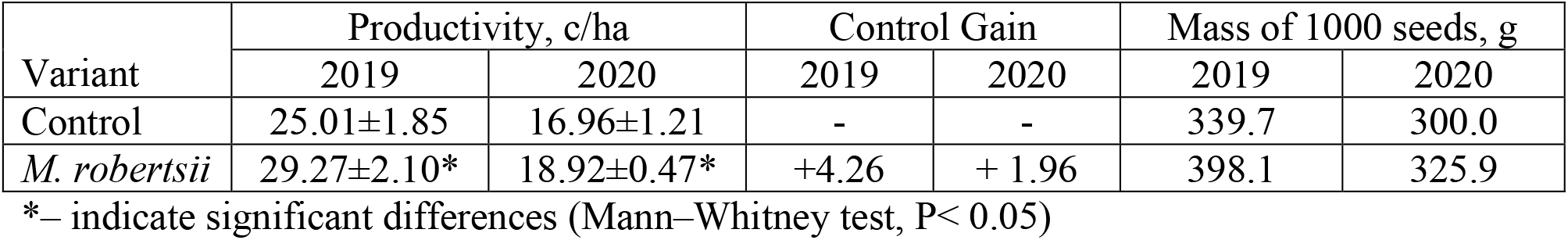
Effect of M. robertsii on bean productivity

## 4. Discussion

As a result of the studies, it was found that the treatment of bean seeds with the entomopathogenic fungus *M. robertsii* stimulated the vegetative development of plants and increased productivity and resistance to phytopathogens. It should be noted that the presented work was carried out in open ground conditions in the territory of western Siberia.

The influence of *Metarhizium* and *Beauveria* fungi on growth stimulation has been previously reported in various plant species, including legumes (Behie *et al*. 2015). Most likely, these effects were observed due to an increase in the supply of nitrogen to plant roots through the mycelium of entomopathogenic fungi and their synthesis of hormone-like compounds and regulatory proteins that changed the metabolism of the plants (Behie and Bidochka, 2014). The interaction of *M. robertsii* with plants can manifest in the form of symbiotic endophytic colonization, which allows obtaining nutrients, protecting the host plant from phytopathogens, and reducing damage from phytophages, which was summarized in a review by Vega (2018) and shown in several later works (St. Leger and Wang, 2020; Barelli *et al*. 2020; Tomilova *et al*. 2020; Hu and Bidochka, 2021).

This study found that seed treatment with *M. robertsii* had a positive effect on aboveground biomass and plant height (an increase of 6-16 cm). Similar data are consistent with the results of other researchers on various plants, including legumes (Khan *et al*. 2012). This is attributed to several reasons: an additional supply of nitrogen to plant roots through the mycelium of endophytic fungi, the synthesis of hormone-like substances by fungi that stimulate plant growth, or the induced production of regulatory proteins that change the metabolic activity of plants (Behie *et al*. 2015). The treated plants had a more developed root system. It was noted that the number of rhizobial nodules was less than that in the control. Similar results were obtained in a study by Barelli and coworkers (2020), which showed that treating beans with the fungus *M. robertsii* increased the number of *Bradyrhizobium* from the group of nitrogen-fixing bacteria and suppressed the development of root rot caused by *F. solani f. sp. phaseoli*, while in the control variant (sterilized soil), symptoms of the disease were visible in the hypocotyl and the upper part of the plant root. Along with this, *M. robertsii* promotes growth and stimulates the proliferation of root hairs on the roots of bean plants (Sasan and Bidochka, 2013); in particular, due to the production of indolyl-3-acetic acid. Further research can focus on quantifying the effect of *M. robertsii* on the root system and the activity of the symbiotic activity of beans.

It has been established that the treatment of faba bean seeds with *M. robertsii* fungus increases the resistance of plants to a complex of diseases. Faba beans treated with the entopathogenic fungus *M. robertsii* were less affected by the complex of diseases. These effects were consistently manifested in two seasons throughout the entire growing season of the plants. In particular, the control plants had lower weight, plant height and yield. This may have been due to both the activation of the immune responses of bean plants under the influence of *M. robertsii* and the direct effect of entomopathogenic fungi on pathogens (Behie *et al*. 2015; Barelli. *et al*. 2020; Barra-Bucarei *et al*. 2019; Hu and Bidochka, 2021).

After the presowing treatment of beans with the entomopathogenic fungus *M. robertsii*, there was a tendency towards a decrease in the level of infection with root rot pathogens. This may have been because the entomopathogenic fungus *M. robertsii* suppressed the germination of pathogens on a nutrient medium. This has been confirmed by a number of studies carried out *in vitro* when studying the antagonistic effect of entomopathogenic fungi on a number of phytopathogens (Sasan and Bidochka, 2013; St. Leger and Wang, 2020).

Five weeks after sowing in 2019 and four weeks after sowing in 2020, in the treated variants, the development and prevalence of root rot caused by a complex of pathogens decreased. The results obtained are in agreement with the studies carried out previously. Thus, a study by Sasan and Bidochka (2013) demonstrated that *M. robertsii*, when studied *in vitro* and in vivo, showed antagonism to the causative agent of bean root rot caused by *Fusarium solani f. sp. phaseoli*. In studies by Ravindran et al. (2014), the metabolite *Metarhizium* ssp. showed inhibitory activity against the phytopathogenic fungi *F. oxysporum, Cladosporium herbarum* and *Curvularia clavata*.

We found that during the entire growing season, there was a significant decrease in the (DDI), with the infection of the aerial organs of bean plants from powdery mildew and chocolate spots, and a tendency towards a decrease in the DDI of *Fusarium*. The ability of *M. robertsii* to act antagonistically against phytopathogens could be related to various complex mechanisms and has not yet been clearly identified (Vega, 2018; Jaber *et al*. 2018; Barra-Bucarei *et al*. 2019). This is due to adaptations such as competition for a niche or resources, antibiosis, parasitism, and induced systemic resistance (Jaber and Ownley, 2017). According to others, this is the stimulation of a plant-induced response or the production of secondary metabolites of *Metarhizium*, which inhibit the growth of phytopathogens (Sasan and Bidochka, 2013; Ravindran *et al*. 2014).

## 5. Conclusion

This work is the first study of the influence of *M. robertsii* entomopathogenic fungi on faba bean grown under field conditions in the continental climate of Siberia. It was found that presowing treatment of seeds reduced the severity of the complex of phytopathogens that were observed on plants during the growing season. In addition, entomopathogenic fungi stimulated the formation of active nodules on the roots, activated the vegetative development of plants and contributed to an increase in productivity. The data obtained represent an essential basis for setting up further detailed studies aimed at improving knowledge about the ecological role of entomopathogens in agroecosystems, as well as issues related to the effective use of entomopathogenic fungi as endophytes in agricultural crops.

## Acknowledgements

The work was done with the support of the Russian Foundation for Basic Research Grant 122011800141-7

